# Workflow and instrumentation for multiscale correlative imaging in voluminous cryogenic specimens

**DOI:** 10.64898/2026.10.14.759080

**Authors:** Edoardo D’Imprima, Evgenia Zagoriy, Saskia Mimietz-Oeckler, Robert Kirmse, Andreas Hallady, Philip Nkemdirim Mbonu, Paolo Ronchi, Martin Jechlinger, Julia Mahamid

## Abstract

Cryo-electron tomography (cryo-ET) enables molecular-resolution imaging of intracellular environments, but is constrained by the need to generate electron-transparent sections of the biological specimen and thus a limited imaging volume. To alleviate such limitations, we present a multiscale correlative workflow that links cryo-confocal fluorescence microscopy (cryo-FLM), cryo-focused ion beam-scanning electron microscopy (cryo-FIB-SEM) volume imaging (cryo-vEM), and cryo-FIB serial lamellae lift-out for cryo-ET in high-pressure frozen voluminous specimens. An integral part of this workflow is a sample holder that preserves specimen orientation across instruments, reduces manual handling, and overcomes stage-tilt limitations in cryo-FIB-SEM hardware. Applied to vitrified colorectal cancer organoids, we demonstrate that this workflow enables fluorescence-guided site preparation for cryo-vEM, which provides biological context beyond the limited field of view of cryo-ET, and targeted lift-out for fabrication of multiple lamellae from defined regions of interest. The reconstructed tomograms reveal subcellular architecture and cell-cell interfaces. This workflow establishes a modular methodological framework for continuum-resolution cryogenic imaging of complex 3D biological specimens.

## Introduction

Cryo-electron tomography (cryo-ET) is transforming structural and cell biology by enabling direct visualization of molecular machines and assemblies in their native cellular environments^1,2^. While early studies were limited to thin specimens such as bacteria^3^, peripheries of adherent cells^4,5^, or isolated organelles^6^, innovations in vitrification^7^ and cryo-focused ion beam (cryo-FIB)^8–13^ micromachining, including lamella lift-out^14,15^ and serial lift-out^16,17^, have extended cryo-ET to whole cells, multicellular specimens and tissues^18–20^, providing unprecedented insight into complex biological systems.

The electron mean free path at 300 keV typically employed for cryo-ET restricts the usable thickness of biological specimens to 200–300 nm^21^. The ~200 nm-thick cryo-FIB lamellae, combined with the limited field of view required for molecular-resolution cryo-ET (up to 1 μm^2^), ultimately define an extremely small imaging volume, with each tomogram capturing only a minute fraction of the biological context. For example, assuming 3000 µm^3^ as an average volume of a HeLa cell^22^, a typical tomogram acquired at 2 Å/pixel covers less than 0.007% of the total cell volume. The severity of this sampling problem scales with specimen complexity^23^. In multicellular specimens, such as small model organisms or tissues, regions of interest are embedded deep within a large vitrified volume, which poses challenges to targeting specific regions of interest and further exacerbates the sampling limitations. Thus, correlative approaches that provide contextual (ultra)structural information over larger sample volumes are needed to bridge this information gap. In particular, serial cryo-focused ion beam-scanning electron microscopy (cryo-FIB-SEM) for volume imaging (cryo-vEM) has emerged as a modality that captures the three-dimensional (3D) ultrastructure of vitrified cells and tissues over volumes of hundreds of cubic micrometers in a label-free manner ^24–26^, providing the multicellular context that individual tomograms lack, albeit at lower resolution. Finally, to harness the full potential of cryo-electron microscopy (EM) imaging modalities that bridge different spatial resolutions in studies of complex biological systems, it is essential to combine molecular specificity with the ultrastructural context. Here, correlative light and electron microscopy (CLEM) joins the complementary strengths of EM and fluorescence light microscopy (FLM), enabling molecular localization within fields of view up to several square millimeters and connecting functional signals with nanometer-scale organization across cells and tissues. CLEM entails diverse elaborate workflows designed to make specimens amenable for both light and electron microscopy, preserving them across multiple imaging modalities^27^. Because biological specimens must be preserved in a near-native, vitrified state for cryo-ET, we focus here on the cryogenic implementation of CLEM (cryo-CLEM), where the challenges of sample preservation, spatial registration, and instrument compatibility differ fundamentally from those at room temperature^2,28–30^.

Solutions for cryo-correlative imaging now exist for individual steps of the workflow, including cryo-confocal FLM^31–34^, cryo-stages for FIB-SEM microscopes cooled either by conduction via metal bands^35^ or cold nitrogen gas through a heat exchanger^8^, and nanomanipulators that operate at liquid nitrogen temperature for lamella lift-out^14,17^. However, comprehensive workflows that ensure compatibility and optimize for imaging quality across all relevant modalities for voluminous high-pressure-frozen specimens are still in early stages of development. Here, we present the development of a custom specimen holder that facilitates transfers between microscopes, addressing limitations associated with cryo-stages cooled via conductive metal bands, and demonstrate a multiscale correlative pipeline that combines cryo-confocal microscopy, cryo-FIB-SEM vEM, and serial lift-out preparation for cryo-ET.

## Results

### Workflow overview

We illustrate an approach that integrates cell biological, ultrastructural, and structural investigation, exemplified on an *in vitro* model of patient-derived colorectal cancer organoids^36^ preserved in a near-native state by high-pressure freezing (HPF)^37^. To improve the yield and reproducibility of organoid preparation from our previous studies^36^, we implemented a high-throughput, low-Matrigel, 3D cell culture method ^38^ (Methods), which yielded >10,000 organoids per plate (Fig. 1A). High-throughput 3D culture is increasingly important for linking phenotypic heterogeneity with structural variability. By integrating such culture systems with targeted cryo-ET, the workflow aims to provide a route toward systematic structural phenotyping of patient-derived material. Immunostaining of key cell polarity markers confirmed preservation of the organoid 3D epithelial architecture under these culture conditions (Fig. 1B). The organoids were harvested (Fig. C) into a tube with optimized culture medium^39^, supplied with cell-permeable live dyes for fluorescent labeling of cellular organelles such as nuclei, mitochondria, and lipid droplets to facilitate light microscopy characterization (Fig. 1D), and subsequently transferred into standard 3 mm-diameter HPF carriers for vitrification (Fig. 1E).

**Figure 1.**
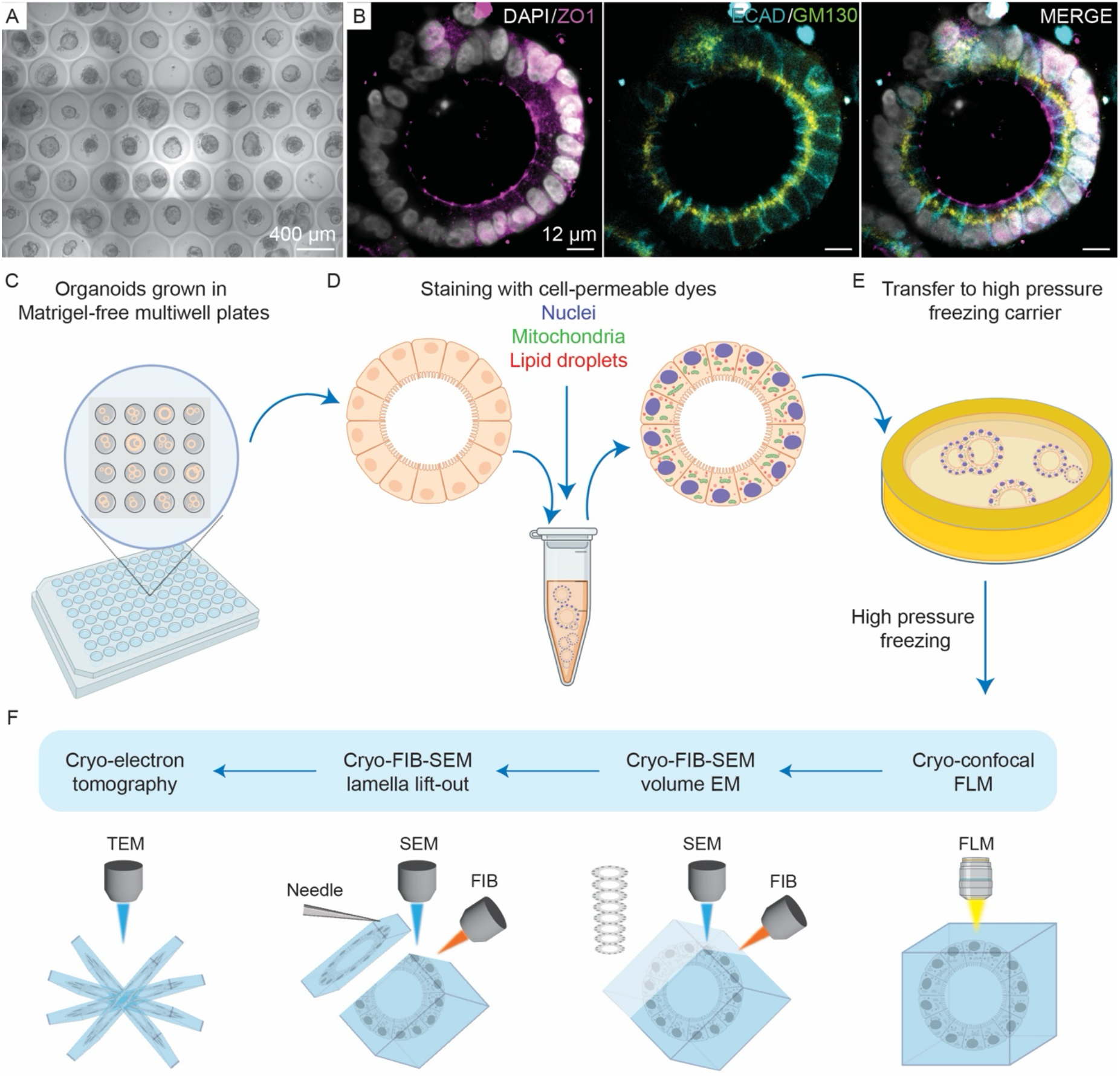
Overview of the multiscale correlative imaging workflow for voluminous cryogenic specimens. (A) Representative bright-field light microscopy image of organoid cell culture in Gri3D microwells^38^. (B) Representative confocal images of individual organoids grown in Gri3D microwells and immunostained for tight junctions (ZO1, magenta), E-cadherin (ECAD, cyan), and Golgi matrix protein 130 (GM130, yellow). Nuclei are stained with DAPI and shown in white. (C-F) Workflow overview of the organoid 3D cell culture in low-Matrigel microplates (C), resuspension of the organoids in medium with organelle-specific live dyes (D), vitrification by HPF (E), and the continuum resolution cryogenic imaging workflow (F).

The next steps constitute a cryogenic continuum-resolution imaging workflow that links fluorescence imaging to cryo-ET via cryo-FIB-SEM vEM and serial lift-out (Fig. 1F): cryo-confocal FLM aims to provide maps of the labeled organelles within complete organoids, which contain critical correlative information that guides the subsequent cryo-EM-based imaging and cryo-FIB micromachining steps. Cryo-vEM on selected organoid regions bridges the fluorescence data with ultrastructural detail, further revealing unlabelled subcellular features. Guided by the information derived from cryo-FLM and cryo-vEM, large volumes from the HPF specimens are extracted by utilizing a cryo-nanomanipulator and following a serial lift-out procedure^17^ to prepare lamellae on an EM grid mounted on the sample holder within the microscope chamber. The cryo-lift-out lamellae are imaged by cryo-ET to obtain 3D molecular-resolution data as the final step of this workflow.

### Hardware overview

In this work, the multiscale cryo-correlative imaging workflow for HPF specimens was developed and implemented using the following hardware setup: a cryo-fluorescent confocal Leica TCS SP8 upright microscope, a Leica cryo loading station (EM VCM), high vacuum coater (Leica EM ACE600), and vacuum cryo-transfer system (EM VCT500). For the FIB-SEM, we equipped a Zeiss Crossbeam XB550, chosen for its demonstrated capability in cryo-FIB-SEM volume imaging^24,40^, with a Leica cryo-stage and shield directly connected via copper bands to the liquid nitrogen (LN_2_) reservoir mounted on the microscope (Fig. 2A, B). In steady-state operation, the cryo-shield and stage are maintained at −190 and −160 °C, respectively, by a Leica auto-refill device with an integrated LN_2_ level sensor, which maintains a constant LN_2_ supply in the reservoir. With its 25 L capacity, the auto-refill system supports up to 14 hours of continuous cooling without user intervention. The Dewar can be refilled during microscope operation, as it is not in direct contact with the LN_2_ reservoir body, allowing for continuous cryogenic operations of up to a week. The software controlling the Leica cooling equipment is integrated with the FIB-SEM microscope, ensuring continuous monitoring and communication between the systems. Prolonged continuous operations were a particular consideration for complex imaging workflows, where experiments on a single sample may require several days.

**Figure 2.**
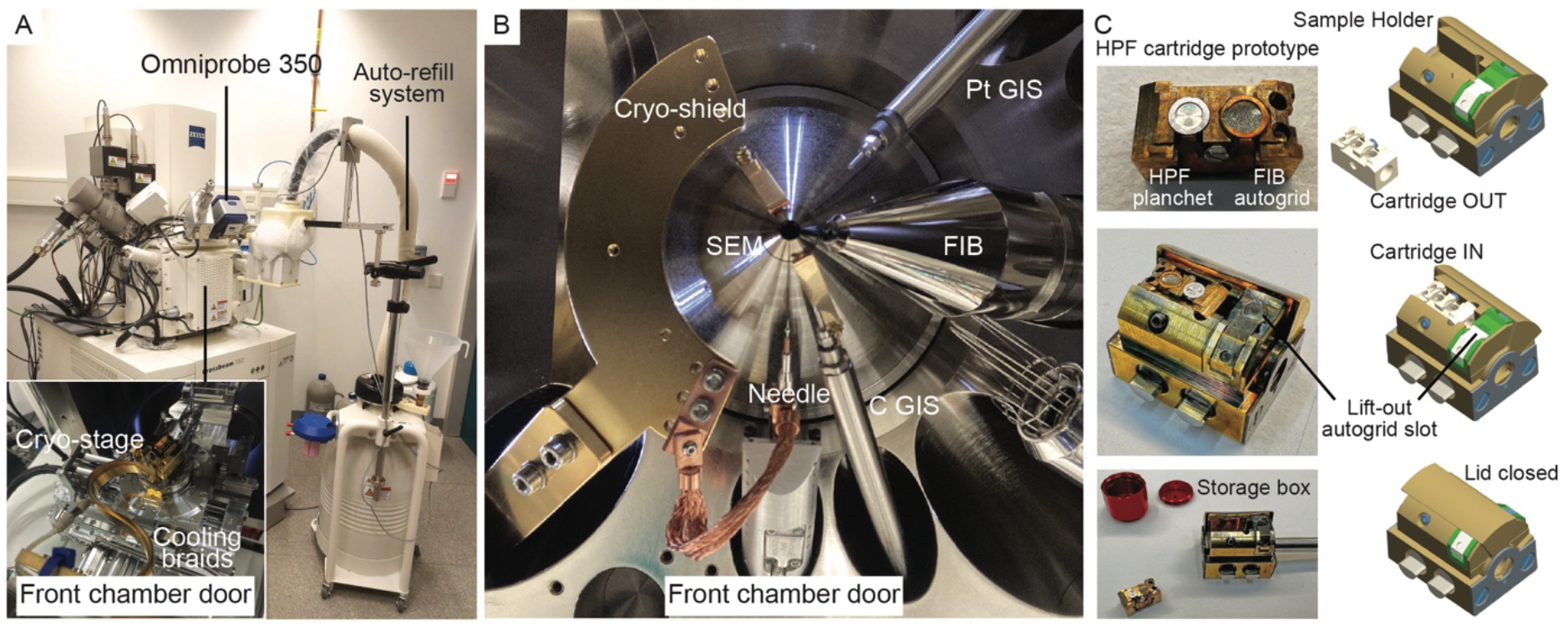
Overview of the hardware employed in this work and the design of a dedicated sample holder. (A) The FIB-SEM microscope equipped with the nanomanipulator (OmniProbe 350 cryo) and the LN_2_ auto-refill system. The inset shows the cryo-stage inside the microscope chamber and its connection to the cooling system via copper bands. (B) View within the microscope chamber towards the pole-piece showing the relative positions of the cryo-shield, nanomanipulator needle, Gas Injection Systems (GIS), SEM, and FIB columns. (C) Images of the developed sample holder (left panel). Top: cryo-FLM cartridge with HPF carrier and autogrid-clipped grid. Two holes on right facilitate handling with tweezers; center: fully assembled sample holder. Position of a recipient grid for lamella from lift-out preparations is indicated; bottom: sample holder connected to the Leica VCT transfer rod with the cryo-FLM cartridge out, and cartridge storage box. 3D rendering of the developed sample holder (right panel). Top: with cartridge out and lid open; center: fully assembled sample holder with cartridge inserted and the lid open; bottom: the same with the lid closed.

The FIB-SEM was additionally equipped with a needle-based cryo-nanomanipulator (Oxford Instruments OmniProbe 350 cryo, Fig. 2B). The nanomanipulator needle (AutoProbe 300/400) connects to the Leica cryo-shield via flexible copper braids that provide a stable temperature of −185 °C, measured at the base of the needle holder (Fig. 2B). The nanomanipulator is controlled by a stand-alone software through a graphical user interface (GUI) that enables motion control along the x, y, and z axes at variable speeds ranging from 50 to 500,000 nm/s. Calibration of the needle tip at the coincidence point of the FIB and SEM beams allows saving multiple positions in the 3D space that can be recalled, facilitating convenient and safe movement from the parking position to a few hundred nanometers from the specimen surface.

### Design of a dedicated sample holder

To streamline transfer and processing of HPF specimens across imaging and micromachining modalities, we designed and prototyped a dedicated holder (Fig. 2C). The design aims to minimize manual handling, simplify image registration by maintaining the same global orientation of the specimen, and accommodate the vastly different geometric requirements of cryo-FIB-SEM vEM imaging, micromachining, lamellae lift-out, and polishing (Fig. 3).

**Figure 3.**
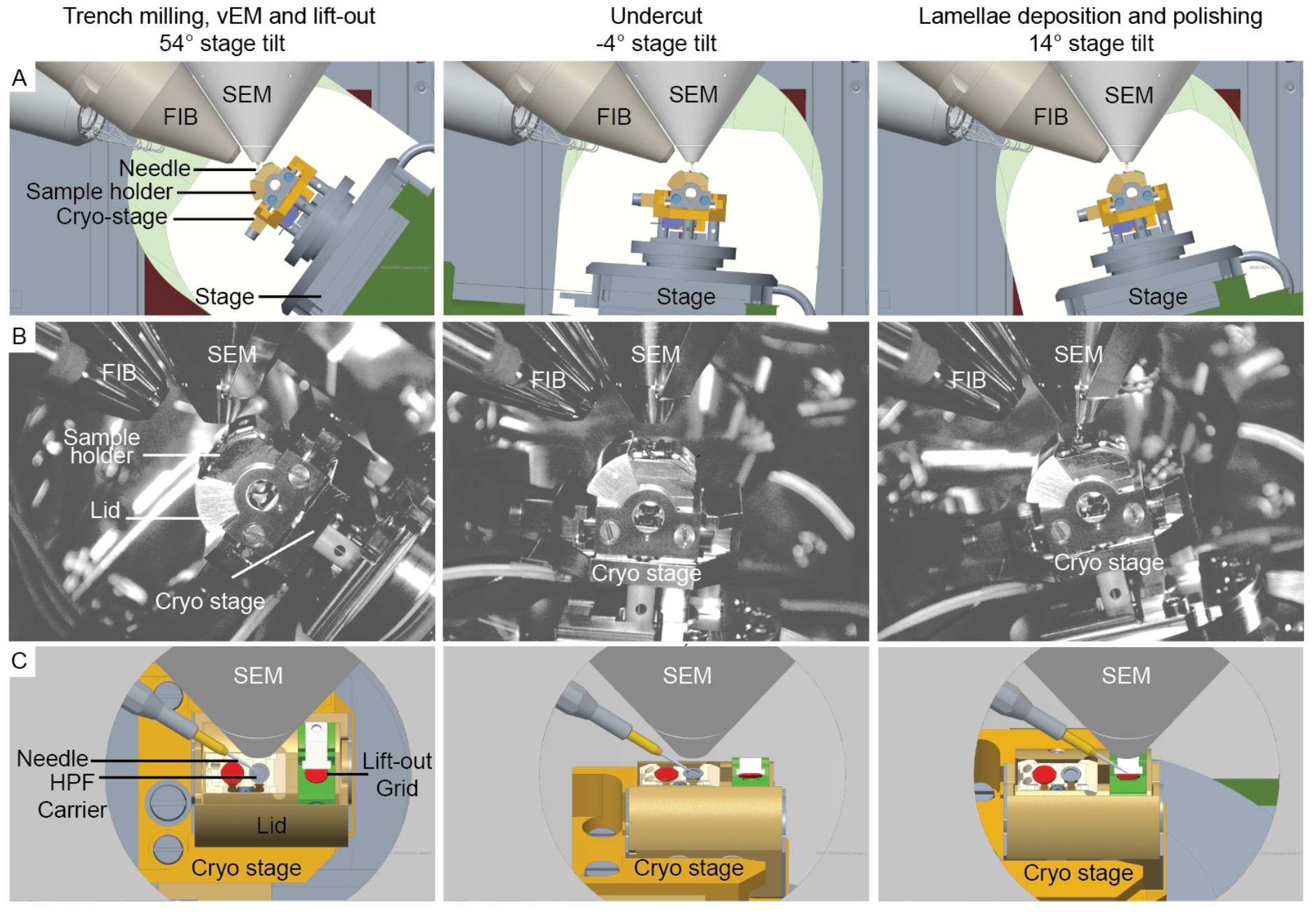
Positioning of the dedicated sample holder within the FIB-SEM chamber required for cryo-vEM imaging and lamellae lift-out operations. (A, B) 3D rendering (A) and CCD camera images (B), viewed from the door of the FIB-SEM chamber. (C) 3D rendering viewed from the FIB column perspective. Color highlights: HPF carrier (grey), autogrid-clipped grids (red), lift-out recipient autogrid clamp (green).

The central component is a cartridge that houses an HPF carrier and has an additional slot that can accommodate an autogrid-clipped EM grid (Fig. 2C), thereby extending the potential application of this setup to “waffle”-type samples^7^. The cartridge is designed based on previous versions that are compatible with imaging in a cryo-FLM equipped with a Leica cryo-stage (Leica TCS cryo-SP8 or STELLARIS 5 Cryo). Both the HPF carriers and autogrid-clipped grids are secured within the cartridge using metal springs. Mounting of the vitrified specimens is performed using a standard and commercially available transfer system. To aid handling, the cartridge features two holes that fit the tips of sharp tweezers. Customized metal boxes enable storage in LN2 and re-use of the cartridges for downstream applications. Following cryo-FLM imaging and selection of a suitable HPF specimen, the cartridge containing the HPF carrier is mounted in the dedicated sample holder, thus retaining the HPF carriers’ orientation in the transfer from cryo-FLM to cryo-FIB-SEM. Mounting of the cartridge into the holder is performed using a Leica cryo loading station (EM VCM). The holder is designed to allow the cartridge to slide directly into its docking position, where it is secured by tightening a metal screw (Fig. 2C). The dedicated sample holder also features a slot pre-tilted by 40° relative to the cartridge, designed to accommodate an autogrid-clipped EM grid that serves as the recipient for the lift-out lamellae. A metal lever secures the recipient grid and locks it in place with a spring mechanism (Fig. 2C). Closing the holder lid provides protection of the cryogenic specimen from environmental ice contamination. Transfer to the FIB-SEM is aided by a vacuum cryo-transfer system (EM VCT500) and the Leica software-controlled load-lock system.

Inside the FIB-SEM microscope, copper bands connecting the cryo-stage to the external LN_2_ reservoir provide stable stage cooling to cryogenic temperatures. However, the copper bands’ length and steric hindrance between the bands and the stage limit the stage rotation to −160° and tilt range from −4° to +54°. Our holder design accounts for these limitations, allowing the execution of critical steps for cryo-vEM and lamella lift-out using only three stage positions that are within the permitted motion range (Fig. 3): because the HPF carrier lies on the sample holder parallel to the stage plane, stage tilt of 54° positions the specimen perpendicular to the FIB column. This position allows trench milling and cryo-FIB-SEM vEM imaging. For lift-out, an undercut to release the targeted specimen block from the HPF carrier is performed at −4° stage tilt. The stage is then returned to 54° to attach the nanomanipulator to the specimen block and to sever the remaining lateral connection of the block to the HPF sample, completing the lift-out. Serial deposition of lamellae from the lift-out specimen block is performed onto the clipped recipient grid that is positioned at 40° tilt relative to the stage, allowing for lamella fabrication, deposition, and polishing at 14° stage tilt, which corresponds to a 10° milling angle between the lamella and FIB source (Fig. 3).

### Application of the workflow to HPF 3D cancer organoids

#### Cryo-FLM and registration for FIB-SEM imaging

Using a Leica TCS cryo-SP8 confocal FLM, we imaged the above-described *in vitro* 3D colorectal cancer organoids stained with fluorescent live dyes and high-pressure frozen in 3 mm diameter, 200 μm-deep HPF carriers (Fig. 4A-C, Methods). The HPF carriers were mounted in the designated cartridge (Fig. 2C). The cryo-FLM setup includes a wide-field light path and a confocal path with independent light sources. Fluorescence data acquisition under cryogenic conditions requires the use of air objectives to preserve the sample’s vitreous state. High numerical aperture (NA), long-working-distance objectives are essential for this purpose, enabling efficient signal collection while decreasing the devitrification risk. The cryo-FLM can accommodate a 50x NA 0.90 WD 0.28 mm air cryo-objective equipped with ceramic housing, providing better thermal isolation, and a 50x NA 0.80 WD 1.04 mm HC PL FLUOTAR air objective. We obtained optimal results when imaging the HPF carriers with the 50x NA 0.80 WD 1.04 mm objective, as it allowed focusing deeper into the 200 μm-thick specimen without the risk of colliding with the sample surface. The confocal light path is equipped with two HyD detectors and 4 laser lines (405 nm, 488 nm, 552 nm, and 638 nm), which we used to detect specific signals from cell nuclei (Hoechst-33342), mitochondria (MitoTracker Green), lipid droplets (BODYPY558/568), and actin (SiR-actin), respectively.

**Figure 4.**
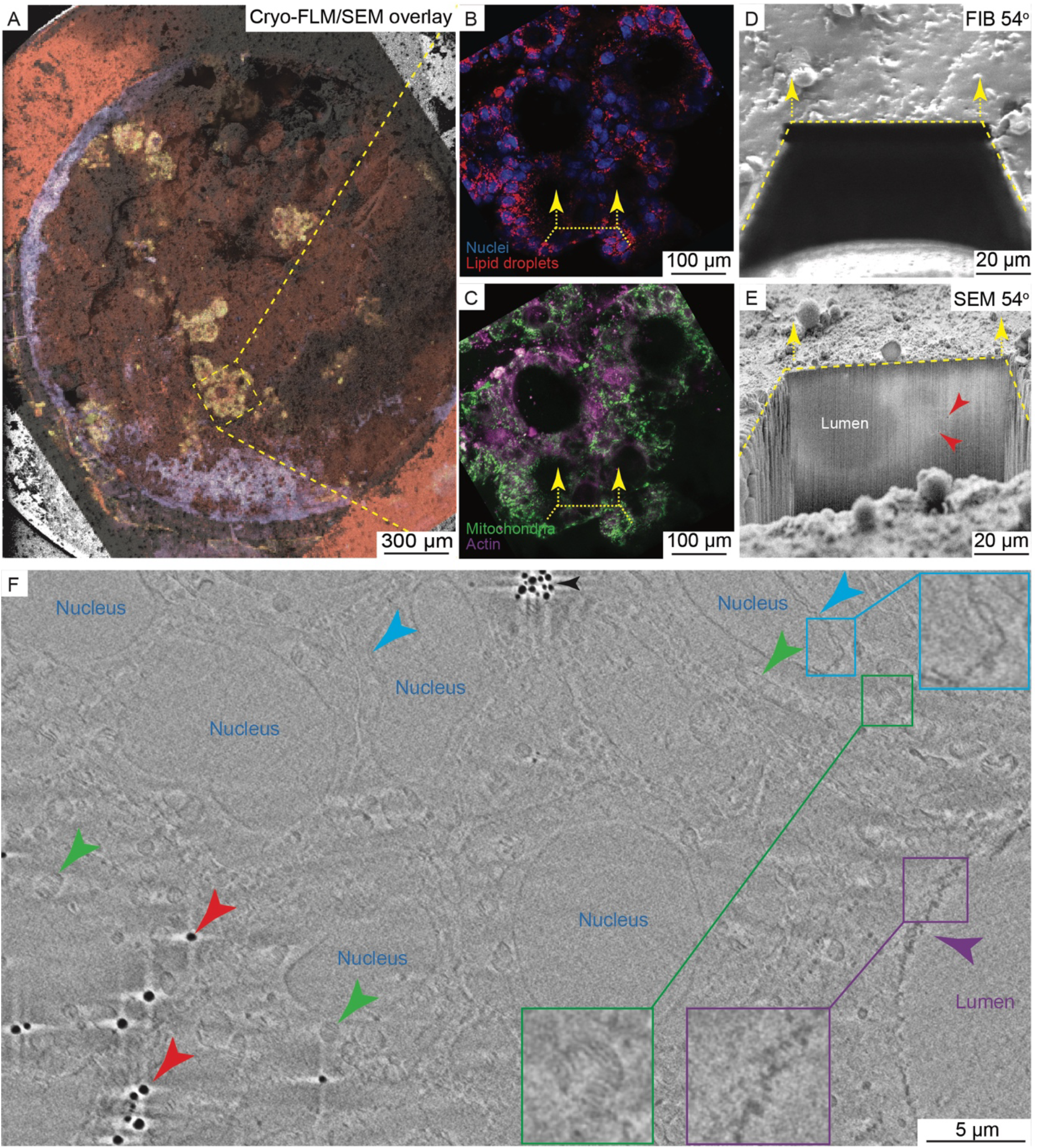
Cryo-FLM-guided cryo-vEM of colorectal cancer organoids. (A) Overview of the fluorescence data registered onto the cryo-SEM image of the HPF carrier’s surface. A cluster of organoids is highlighted and enlarged in (B) blue: nuclei, red: lipid droplets, and (C) green: mitochondria, magenta: actin. Channels are separated for better visualization. Yellow dashed lines (B-E) outline the cryo-FIB trench location (D), and arrows indicate the direction of iterative ablation in the process of the vEM acquisition (E). (D) FIB and (E) SEM views of the opened trench before cryo-vEM. The SEM view allows identification of sample features. Indicated are the organoid lumen and lipid droplets (red arrowheads). (F) Frame from the cryo-vEM data showing cell nuclei, nuclear envelopes (blue arrowheads and inset), lipid droplets (red arrowheads), mitochondria with cristae (green arrowheads and inset), organoid lumen, and microvilli (magenta arrowheads and inset).

The fluorescence data of the HFP sample guided subsequent cryo-EM operations. For that, the cartridge was mounted into the dedicated sample holder within the Leica VCM, the biological specimen was rendered conductive by applying a 40 nm thick layer of metallic platinum (Pt) using a Leica ACE 600 sputter coater (Methods), and the whole assembly was transferred into the FIB-SEM microscope with the aid of the Leica VCT. Image registration onto the SEM image of the HPF carrier acquired at stage tilt 0° was performed within SmartSEM software (Fig. 4A-C) based on surface features that were recognizable in both modalities. This allowed locating and targeting organoids of interest within the bulk of the HPF carrier.

**Video 1. Cryo-vEM of colorectal cancer organoids**. Slicing through the raw and processed data (starting at 00:17) of a volume of 60 × 40 × 10 µm containing parts of 8 cells, showing lipid droplets, mitochondria, cell nuclei, and organoid lumina lined by microvilli.

#### Cryo-FIB-SEM volume imaging

We next performed cryo-FIB-SEM volume imaging to obtain 3D ultrastructural data of the multicellular organoid context, while approaching the portion of the specimen to be subsequently probed by cryo-ET. Prior to the cryo-FIB micromachining, we applied a 2-3 µm-thick organo-metallic Pt layer using the in-chamber GIS at a stage tilt of 0° to shield the underlying beam-sensitive biological material and minimize curtaining during FIB milling (Methods). The correlated cryo-FLM map was used to navigate the specimen and position the target area at the coincidence point of the FIB and SEM beams (Fig. 4A-C). We then tilted the stage to 54° such that the HPF carrier surface was perpendicular to the FIB column, and micromachined a trench of 80 × 80 µm using FIB currents of 7-15 nA, followed by polishing the exposed surface at 1.5 nA (Methods). The polished cross-section showed the presence of biological material (Fig. 4D, E, Methods), revealing characteristic features, such as the outline of the organoids’ lumen and high-contrast lipid droplets. We next performed cryo-vEM imaging using a pixel size of 20 × 20 nm and a milling step of 40 nm at a FIB current of 0.7 nA (Methods). The automated iterative material ablation with the FIB and image acquisition with the SEM were controlled by Atlas 5 software (Atlas Engine using the Nanotomography workflow). As previously reported, vitreous samples suffer from low conductivity and tend to accumulate charge during the SEM imaging, which deteriorates image quality by introducing charging artifacts^25^. To alleviate the charging artifacts, we used a short point dwell time (0.1 msec) of the SEM (2.3 keV, 20 pA), combined with line averaging of 150, which allowed better charge dissipation and, consequently, more consistent contrast in the acquired images. The cryo-SEM images additionally exhibited low signal-to-noise ratio, contrast gradients across the image, curtaining, and drift. To compensate for the drift, we aligned the individual frames with linear stack alignment using Scale-Invariant Feature Transform (SIFT)^41^ (Methods). To improve image quality, we used the VSNR_V2 Fiji plugin^42^ to remove structured and unstructured noise. Finally, we used Contrast Limited Adaptive Histogram Equalization (CLAHE)^43^ to enhance details that were obscured by the large variations in brightness and contrast in the images (Methods). The processed data captured a volume of 60 × 40 × 10 µm, containing parts of 8 cells, and clearly showed lipid droplets, mitochondria with discernible cristae, cell nuclei delineated by nuclear envelopes, and organoid lumina lined by microvilli (Fig. 4F, Video 1).

#### Serial lift-out and cryo-ET

Having acquired ultrastructural cryo-vEM data, we proceeded to generate lamellae for molecular-resolution cryo-ET from the same specimen area via a serial lift-out protocol^17^ with some modifications (Methods). As a first step, we used a 54° stage tilt and FIB currents of 7-15 nA to shape the sample block by opening trenches at the sides and the back of the targeted area (Fig. 5A, B). We next introduced an undercut at −4° stage tilt and 7 nA FIB current, thereby detaching the bottom of the block from the rest of the HPF specimen. The resulting block remained attached to the bulk specimen by material on the side. After returning to a 54° stage tilt, we inserted the cryo-nanomanipulator needle and attached it to the sample block (Fig. 5C): we exploited the redeposition of sputtered materials for the welding of the nanomanipulator to the sample block^16,17^. Following current common practice, we micromachined a cube of copper from standard EM grids and welded it to the tip of the tungsten needle of the nanomanipulator ahead of the cryo-FIB-SEM session. This copper block then serves as the interface between the needle and the lift-out target and provides the material for their attachment (Methods). Following welding of the block, we cut off the remaining connection to the HPF specimen, lifted the sample block from the trench (Fig. 5D), and moved it to the receiving EM grid mounted at 10° relative to the block plane (Fig. 5E). There, the sample block was positioned such that its edges rested on the grid bars, from which material was used to weld the bottom of the sample to the grid by redeposition (Fig. 5F). A lamella was then cut off, and the block moved onto the next grid position for the generation of additional lamellae by repeating the process (Fig. 5G). In this manner, we generated 4 × 5 µm thick lamellae from one sample block, which were then thinned down to 0.6-0.8 µm and finally polished to the target thickness of 200 nm (Fig. 5 H, I, Methods). The recipient grid with the resulting lamellae was transferred into a cryo-TEM, where lamella maps were acquired at intermediate magnification (11.09 Å/pixel) (Fig. 5 J) to assess ice quality and select areas for high-resolution cryo-ET data acquisition (3.03 Å/pixel) (Fig. 5 K, L). The reconstructed tomographic volumes showed subcellular and molecular detail, as well as views of cell-cell interfaces within the multicellular specimen.

**Figure 5.**
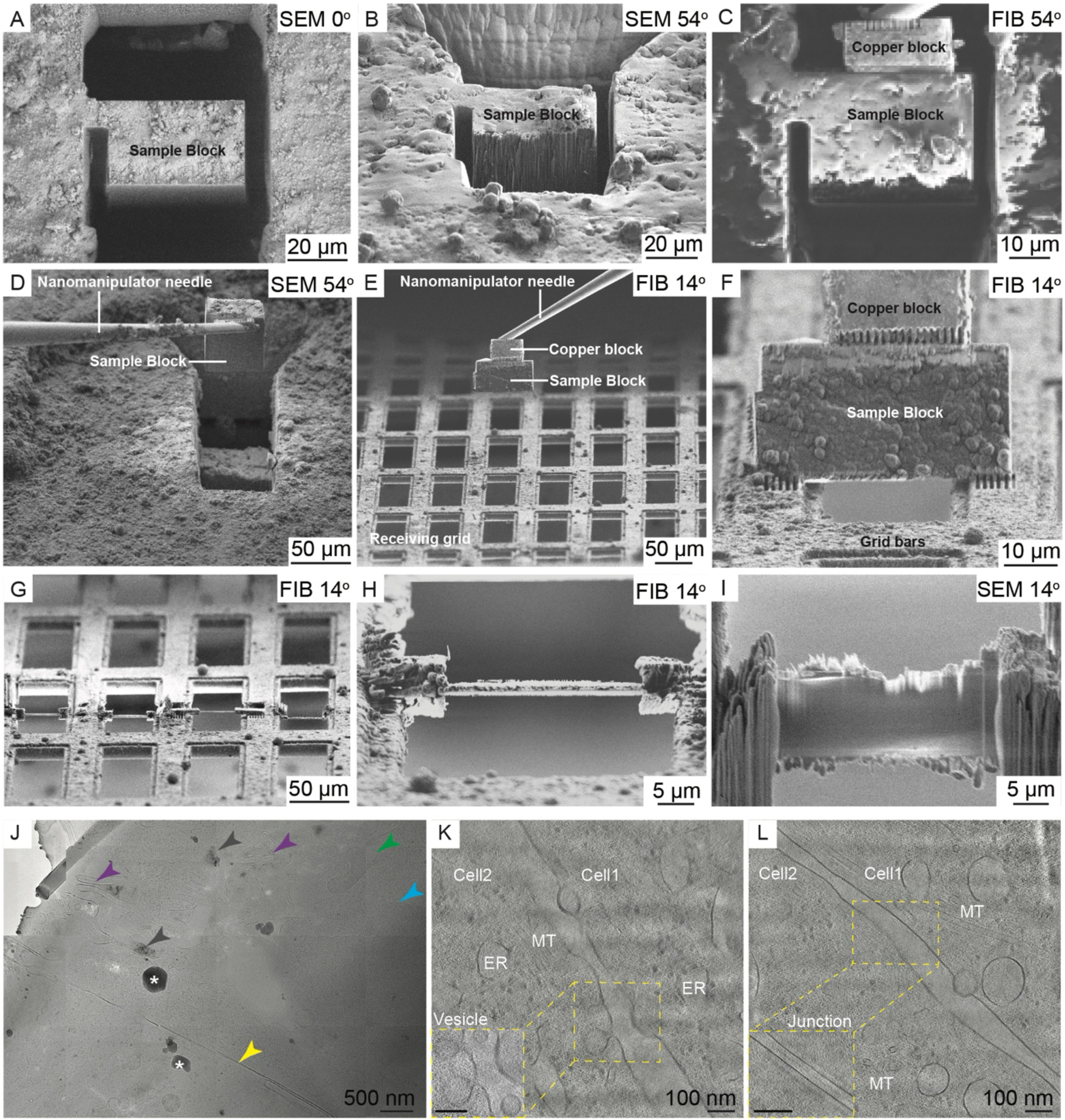
Targeted serial lift-out and cryo-ET of colorectal cancer organoids. (A) SEM view at 0° stage tilt of the sample block for lift-out. (B) SEM view of the same specimen location at a 54° stage tilt exposes the side of the sample block onto which the nanomanipulator needle will be attached. (C) The nanomanipulator and the sample block are connected via the copper block; an image scan rotation of 180° is applied to aid perception. (D) Lift-out of the sample block from the trench. Orientation similar to that shown in B. (E) Transfer of the lift-out sample block to the receiving EM grid (Cu 100 × 400 mesh). (F) Welding of the sample block to the receiving grid using copper redeposition from the grid bars. (G) Consecutive slicing of 5 µm-thick lamellae off the block and welding to the grid bars. (H, I) Rough milled lamella viewed from the FIB (H) and SEM (I) perspective. (J) Cryo-transmission electron micrograph of the polished lamella. Indicated are crystalline ice reflections in the extracellular matrix (grey arrowheads), ice contaminants introduced during transfers (*), microvilli (magenta arrowheads), mitochondrion (green arrowhead), nuclear double membrane (blue arrowhead), and cell-cell interface (yellow arrowhead). (K, L) Representative tomographic slices showing the cell-cell interface with endoplasmic reticulum (ER), microtubules (MT), and vesicles (K) or the location of a putative tight junction^44^ (L). Insets in the bottom-left corners show different tomographic slices within the same cryo-ET reconstructions.

## Discussion

This work establishes a cryogenic multiscale correlative imaging workflow and primarily addresses technical limitations associated with cryo-stages cooled via thermal conductivity of metal bands, which impose strict restrictions on stage movements within FIB-SEM instruments. Such cryo-stages are currently employed in cryo-FIB-SEM microscopes across various commercial platforms, including Zeiss^45^, Tescan^46,^ and Jeol^47^. The herein-developed sample holder and workflow provide a universal solution for such hardware, improving the availability of cryo-sample imaging and preparation approaches for in-cell structural biology studies across research laboratories and institutions.

By combining high-pressure freezing, fluorescence-guided navigation, contextual cryo-vEM, and multi-lamella extraction, this workflow addresses two current constraints in cellular cryo-ET: complementing the scarce imaging volume per tomogram and facilitating targeting of specific regions within voluminous, heterogeneous multicellular specimens. Our multiscale workflow expands the framework of “continuum-resolution imaging,” in which fluorescence and ultrastructure are not simply correlated retrospectively, but used in sequence to direct destructive imaging and preparation steps. In contrast to workflows that rely exclusively on fluorescence overlays for FIB targeting, the inclusion of cryo-vEM introduces an intermediate validation layer that can be advantageous in complex 3D multicellular systems.

### Universal sample holder enables robust integration across modalities

Cryogenic CLEM workflows often suffer from geometric incompatibilities between instruments and limited stage motion due to cooling hardware. The sample holder developed here preserves specimen orientation across cryo-FLM, cryo-vEM, and lamella lift-out preparations. By constraining the workflow to three optimized stage configurations within the cryo-FIB-SEM microscope, our design circumvents rotation and tilt restrictions imposed by the conductive cooling bands while maintaining compatibility with serial lift-out procedures. The compatibility of serial lift-out with HPF-preserved 3D cultures is consistent with previous demonstrations in whole organisms and tissues^14,16–18,48,49^, reinforcing its broad applicability.

Such hardware-level integration facilitates reproducibility and automation^12^. The ability to transfer HPF carriers without reorientation reduces registration error and minimizes manual handling, which represents a significant source of sample loss and ice contamination in cryogenic workflows. The design also supports potential adaptation to waffle-based approaches for grid-mounted specimens^7^, increasing its versatility.

### >Multiscale correlation to overcome the field of view disparity

Cryo-ET enables molecular-resolution imaging of intact cells, but is intrinsically restricted to lamellae up to 300 nm thickness. Advances in cryo-FIB milling and lift-out have further expanded cryo-ET from studies of single cells to whole organisms and tissues, including pathological human samples^14,16,49^. Yet, retaining and connecting contextual information across spatial scales in such complex 3D biological systems remains a challenge.

Our workflow reduces data registration uncertainty at each imaging scale. Cryo-FLM places molecular markers throughout the HPF carrier, providing specificity that is otherwise absent from the cryo-FIB-SEM vEM and cryo-ET data, where contrast originates only from structural properties of subcellular features. Our workflow uses a cryo-confocal fluorescence imaging modality. Although this introduces technical hurdles associated with specimen transfers between microscopes in comparison to wide-field systems integrated into the FIB-SEM microscope chamber^50,51^, it provides superior resolution (especially in the z dimension), better spectral separation, and reduced out-of-focus background. Cryo-vEM then offers valuable ultrastructural-resolution context in heterogeneous specimens across hundreds of cubic micrometers, and can facilitate 3D localization of cellular structures^24,40,52^. While cryo-vEM does not achieve molecular resolution, it expands the contextual volume relative to cryo-ET and provides spatial continuity across extraction sites.

### Targeting regions of interest in cryogenic specimens

For simple specimen geometries, such as bacterial, yeast, or single mammalian cells plunge-frozen on EM grids, direct cryo-FLM targeting followed by on-grid cryo-FIB milling is typically sufficient: regions of interest are close to the sample surface, thereby reducing optical aberrations ^30^; correlation can be made straightforward by introduction of fiducials; and throughput (lamellae/day) is maximized^8,12,30,31,53^. Our workflow is advantageous when these assumptions break down: in thick HPF carriers containing heterogeneous organoids or tissue specimens, the targets are buried at variable depth, and surface landmarks detectable across modalities are limited. In this regime, the intermediate cryo-vEM step can provide structural validation for the registration accuracy of the cryo-FLM to the cryo-FIB-SEM before the destructive sample block preparation step and enables rational serial lift-out to generate multiple lamellae sampled across a defined 3D region, potentially improving targeting confidence and yield.

### Limitations

Despite the benefits of the presented workflow, several limitations persist. HPF does not reliably achieve complete vitrification of heterogeneous voluminous samples, resulting in crystalline and thus structurally unpreserved regions. Furthermore, surface ice contamination following HPF and due to transfer steps across microscopes remains a challenge. Improvements in environmental control and transfer systems may help to reduce these effects^10^. Optical distortions and registration errors in fluorescence imaging of thick HPF samples limit the accuracy of correlations in cryo-CLEM^30^. Further developments in cryogenic deep fluorescence imaging approaches may enhance localization precision. Although the workflow presented here improves the overall efficiency of targeted lamella preparation, automation of lift-out preparations is still critically missing. The steps of cryo-FIB-SEM vEM data acquisition, image registration, and data processing remain time-intensive. Dedicated adaptations to improve cryo-SEM imaging of vitreous material where low contrast, charging, and beam sensitivity dominate, through dose-efficient acquisition that integrates multiple short-dwell-time frames^47^, low-voltage native-contrast imaging^54^, and deep-learning-guided data-aware acquisition, along with on-the-fly correlation across imaging modalities^52^, are of key importance to improve scalability.

### Outlook

The growing body of cryo-ET applications to disease-relevant human samples underscores the need for robust and transferable multiscale imaging workflows^28^. The approach presented here integrates specimen preparation, hardware design, and multiscale imaging, with the potential to target specific regions of interest, into a coherent framework compatible with complex 3D specimens. This timing is particularly apt: microwell-based organoid platforms now enable scalable and standardized cohorts, while in cryo-ET the bottleneck has shifted from imaging^55^ to reliable sampling and targeting within thick, heterogeneous specimens. By combining fluorescence specificity, cryo-vEM ultrastructural context, and serial lamella extraction, our workflow enables multi-lamella sampling across phenotypic heterogeneity from a single HPF specimen, and provides a practical path to comparative studies across genotypes and perturbations. In doing so, it narrows the gap between phenotypic cell biology and molecular-resolution analyses by anchoring ultrastructural signatures to functional states in native contexts.

## Materials and Methods

### Human Organoid Culture

HT6 organoids were obtained and maintained as described previously^36^ in Sato medium^39^. The Patient-derived material was used following all required ethics approvals as previously detailed^36^. For the high-throughput culture method adopted here, organoids were grown in anti-adherent Gri3D^®^ 96-well plates equipped with an imaging bottom (SUN Biosciences)^38^. Each of the 96 wells contains 121 microcavities of 400 µm diameter, with each cavity compatible with growth of a single organoid. To generate the cell suspension for seeding, organoids grown in Matrigel domes were extracted using ice-cold basal medium. Briefly, basal medium (DMEM 1x 4.5 g/L D-glucose FluoroBrite medium (Gibco, Cat.# A1896701) supplemented with 10% inactivated FBS (Gibco, Cat.# A5670701), 1% HEPES 1 M, 1% Sodium Pyruvate 100 mM (Gibco, Cat.# 11360070), 1% MEM NEAA 100X (Gibco, Cat.# 11140050), 1% L-Glutamine 200mM (Gibco, Cat.# 25030081), 1% Penicillin Streptomycin solution) was cooled on ice. The medium was kept cold and without FBS to maintain the Matrigel in a liquid state and facilitate subsequent cell dissociation. First, the Sato medium was aspirated without disturbing the Matrigel domes containing the organoids, and 1 ml ice-cold basal medium was added to each dome. 1 ml pipette tips were pre-wetted in ice-cold basal medium and used to scrape the Matrigel domes from the plate bottom. Pipetting 2-3 times disrupted the domes. The suspension was collected into 15 ml tubes, pre-cooled on ice, and topped with an additional 1 ml of basal medium used to rinse the remains from the plates. Up to 4 domes were pooled in each 15 ml tube. The tubes were centrifuged twice at 200 g for 4 minutes at 4° C, the supernatant was discarded, and the pellet was transferred to a new tube pre-wetted with basal medium. To dissociate organoids into single cells, the dissociation medium was added (TrypLE (Gibco™, Cat.#12604039) with 10 µM ROCK inhibitor Y-27632 (Stemcells Technologies Cat. # 72304)), and the tubes were incubated at 37 °C for 3-7 minutes. To time the dissociation, ~100 µl of the organoid dissociation mixture was transferred to a 200 µl tube to directly monitor the dissociation process every minute at an inverted microscope (Olympus CKX41) with CellSence Standard software. When required, to aid dissociation, the mixture was resuspended using P1000 tips pre-wetted in dissociation medium. The dissociation was halted by adding 4 times the volume of fresh basal medium supplemented with FBS to inactivate TrypLE. The cells were then centrifuged at 1,000 g for 5 minutes, the supernatant discarded, and the pellet resuspended in 100 µl fresh Sato medium supplemented with 4% Matrigel. Cell concentration was estimated using a cell counter (Bio-Rad TC20 Automated Cell Counter Cat.#1450102), mixing trypan blue and cell suspension 1:1, and adjusted with additional Sato medium with 4% Matrigel to 1.5-3.5 × 10^5^ cells/ml, corresponding to roughly 50-150 cells per 400 µm microcavity. Cells were then aliquoted into the Gri3D^®^ plates and incubated at 37° C and 5% CO_2_. The 3D cell culture was monitored daily until fully grown organoids were obtained, typically at 10 days post-seeding.

### Immunofluorescent staining of epithelial polarity markers

Immunofluorescence staining (Figure 1) was carried out as previously described with minor modifications to validate the organoids’ epithelial polarity^56,57^. The HT6 organoids were gently washed 3 times with 1x PBS in the Gri3D 96-well plates via the lateral pipetting port to remove the Matrigel. Organoids were fixed in 4% paraformaldehyde (Biozol, Cat.#BOT-22023) for 30 minutes at room temperature without agitation. Post-fixation, organoids were washed 3 times with 1x PBS for 10 minutes each time and permeabilized using freshly prepared 1x IF buffer (pH 7.4), also 3 times for 10 minutes, at room temperature. The IF buffer contained 7.6 mg/ml NaCl (Sigma-Aldrich, Cat.#1.06404), Tween-20 (Sigma-Aldrich, Cat.#P9416), 1.87 mg/ml Na_2_HPO_4_*7H_2_O (Sigma-Aldrich, Cat.#S9390), 0.4 mg/ml NaH_2_PO_4_*H_2_O (Sigma-Aldrich, Cat.#S9638), 0.5 mg/ml NaN_3_ (Sigma-Aldrich, Cat.#S8032), 10 mg/ml BSA (Sigma-Aldrich, Cat.#A9418), 20 µl/ml Triton X-100 (Sigma-Aldrich, Cat.#T8787) at final concentrations. Permeabilized organoids were blocked with 10% goat serum (Jackson Immunoresearch, Cat.# 005-000-121) for 4 hours at 4 °C. Primary antibodies (Mouse-anti GM130 1:100 (BDbiosciences, Cat.# 610823), Rat anti-E-Cadherin 1:150 (Thermo Fisher Scientific, Cat.# 13-1900), Rabbit polyclonal anti-ZO1 1:500 (Thermo Fischer Scientific, Cat.# 61-7300)) were diluted in the primary blocking buffer (5% goat serum in IF buffer) and incubated overnight at 4°C. Organoids were washed the next day with 1x IF buffer 3 times, 10 minutes each, and incubated overnight at 4 °C with secondary antibodies (Goat anti-rat IgG (H + L) Alexa Fluor 647 1:500 (Thermo Fischer Scientific, Cat.# A21247), Goat anti-mouse IgG (H + L) Alexa Fluor 568 1:500 (Thermo Fischer Scientific, Cat.# A11031), Goat anti-Rabbit IgG (H+L), Highly Cross-Adsorbed Alexa Fluor 488 1:500 (Thermo Fischer Scientific, Cat.# A11034)) and 4’, 6’-diamino-2-phenylindole (DAPI) 1 µg/ml (Thermo Fisher Scientific, Cat.# 62248), all diluted in the primary blocking buffer. Finally, the organoids were washed with 90-120 µl of 1x PBS 4 times, 1 hour each, via the lateral pipetting port and mounted with 50 µl of Vectashield Hardset Antifade mounting medium (Biozol, Cat.# VEC-H-1500). After polymerization of the mounting medium, organoids were imaged on a Nikon A1R HD25 confocal microscope (CFI Apo LWD Lambda-S 40x/1.15 WI 0.60 WD 610 µm), at a z-step of 0.5 µm, using the following excitation wavelengths: 405 nm, 488 nm, 561 nm, 640 nm.

### High pressure freezing

We followed the procedure described in Ronchi et al. 2021^58^ with minor modifications. Briefly, to resuspend organoids grown in the Gri3D^®^ plates, the medium was gently pipetted using a P200 pipette fitted with 200 µl tips. The narrow ends of the tips were cut to widen the opening and prevent mechanical damage to the organoids. To prevent organoids from sticking to pipette tips, the tips were pre-wetted in 2% BSA in PBS (Sigma-Aldrich, Cat# A8806). The resuspended organoids were collected in a 15 ml Falcon tube and allowed to gently pellet by gravity in an incubator at 37°C and 5% CO_2_ for 15 minutes. Harvested organoids were stained before freezing for 10-20 minutes in culture medium containing the following dyes in their final concentrations: Hoechst-33342 10 µM (Thermo Fisher Scientific, Cat.# H1399), MitoTracker® Green FM dye 1 µM (Thermo Fisher Scientific, Cat.# M7514), BODIPY 558/568 5 µM (Thermo Fisher Scientific, Cat.# D3922), and SiR-Actin 2 µM (Spirochrome, Cat.# SC001). Finally, 1-2 µl of organoid suspension was pipetted into 3 mm diameter, 200 µm deep aluminium HPF carriers (Wohlwend GmbH, article 241), closed in B-type HPF carriers and high-pressure frozen in an HPM 010 (AbraFluid) or Leica EM ICE. The B-type carriers used for the HPF “sandwich” assembly were first coated with 2% Phosphatidylcholine (Merck, Cat.#61755) in Ethanol (Merck Millipore, Cat.#107017) and allowed to dry to facilitate disassembly post-freezing.

### Cryo-Confocal microscopy

Confocal stack acquisition under cryo-conditions was carried out with a Leica TCS SP8 upright microscope (Leica Microsystems), controlled by Leica Application Suite X 3.5.5.19976 software. The microscope was equipped with a cryo-stage, an HC PL FLUOTAR 50x NA 0.80 WD 1.04 mm DRY objective, and a Leica DFC365 FX camera. The high-pressure frozen organoids in HPF carriers were inserted into the cartridge prototype with the sample side facing the objective. All sample loading and transfer operations were conducted under cryo-conditions, in the LN_2_ vapor phase, using a dedicated loading/transfer shuttle (Leica Microsystems). The loading and transfer steps were carried out in a humidity-controlled room (10-20%) to minimize ice contamination on the sample. After the cartridge was deposited on the microscope stage pre-cooled to −195 °C, we determined the approximate z-position of the sample surface in wide-field illumination mode. Using the Navigator module, we acquired 100 μm deep (steps of 2 μm) z-stack tiles (x/y with 20% overlap) to produce a stitched 3D overview image of the whole carrier at the desired wavelengths. This helped to determine regions of interest for subsequent confocal acquisitions. Confocal stacks were acquired in four channels in two sequences: 405 nm at 5% power Diode laser (HyD1 detector) and 552 nm at 1% power OPSL laser (HyD 3 detector), followed by 488 nm OPSL laser at 1% (HyD1 detector) and 638 nm Diode laser at 1% power (HyD 3 detector), and using the following parameters: pinhole 1 AU, zoom 1, pixel format 1024 × 1024, pixel size in x,y = 0.303 μm, in z = 0.5 μm, bidirectional scan at 600 Hz, z-stacks of 50 μm depth. Confocal stacks were processed with the Lightning deconvolution software module (Leica Application Suite X 3.5.5.19976) at default settings and a number of iterations set to 5 for all channels. After the acquisition, the HPF carriers were retrieved under cryo-conditions for further analysis and stored within the cartridge prototype in the storage box.

### Cryo-FIB-SEM site preparation, imaging and vEM data processing

The cartridge housing the HPF sample carrier was loaded onto the newly developed specimen holder in a LN_2_ bath using a VCM-VCT transfer system (Leica Microsystems). The whole assembly was transferred into the EM ACE 600 (Leica Microsystems), pre-cooled to −170 °C, and sputter-coated with 40 nm Pt. Subsequently, the holder was loaded into a Zeiss Crossbeam 550 via the Leica VCT500 transfer system. Organometallic Pt was deposited using the in-chamber GIS at a stage tilt of 0°. All FIB operations were done at an acceleration voltage of 30 keV. Large trenches were milled with 7-15 nA FIB current, and the cross-section surface was polished with 700 pA or 1.5 nA currents. Precise milling during the volume acquisition was achieved with a current of 700 pA. The SEM imaging was done with an acceleration voltage of 2.3 keV and a current of 20 pA, using an InLens secondary electron detector. Data were acquired at 10 × 10 × 30 nm or 20 × 20 × 40 nm (x,y,z) sampling. To reduce the effects of charging on SEM image quality, pixel dwell time was set to 0.1-0.2 μs and line averaging to 120-150.

Alignment of the acquired FIB-SEM stacks was performed using scale-invariant feature transform (SIFT)^41^ implemented in Fiji^59^. The expected transformation was set to translation, with an initial Gaussian blur set to 1.6 px. T aid data visualization, we used VSNR_V2^42^ (Fiji/ImageJ Plugin) to remove vertical stripe artifacts originating from curtaining with the following settings: a Gabor filter (Ex=1, Ey=1400) and DenoiseH1 algorithm (noise level 0.2-0.3). We repeated the procedure for horizontal lines originating from charging with the Gabor filter, and DH1 denoising set to: Ex=500, Ey=50, noise level 0.3-0.5. White noise was reduced by applying a Dirac filter (noise level 0.2) and a Gaussian blur (sigma 0.75). After VSNR_V2, we used a Contrast Limited Adaptive Histogram Equalization (CLAHE)^43^ to mitigate the effects of spatially varying illumination and high dynamic range intensity distributions (maximum slope set to 1.5). Three cryo-vEM datasets representative of the opening of an organoid lumen (Vol1: 20×20×40 nm sampling rate in x,y,z; total volume ~24.300 µm^3^), the bulk portion of one organoid (Vol2: 20×20×40 nm sampling rate in x,y,z; total volume ~ 7.000 µm^3^), and the tip of an organoid crypt (Vol3: 10×10×30 nm sampling rate in x,y,z; total volume ~80.440 µm^3^) will be deposited on EMPIAR.

### Serial cryo-lift-out

To prepare for the cryo-lift-out procedure, we first clipped a rectangular pattern 100 × 400 mesh copper TEM grid (Agar Scientific, Cat.# AGG2140C) into an autogrid (Gewo Feinmechanik GmbH, Cat.# Q1192OHNE) and loaded it into the newly developed specimen holder in the lift-out autogrid slot. The whole assembly was loaded into the Zeiss Crossbeam 550 at room temperature, rotated 180° with respect to the loading position intended for the main cryo-workflow. This allowed us to overcome the limitation imposed by the stage cooling bands on the rotation range, and to position the TEM grid surface perpendicular to the FIB. We then prepared a 20 × 20 μm copper block out of one of the grid bars using a FIB current of 7 nA. This was achieved by placing rectangular patterns of 22 × 2 μm on 3 sides around the block and 15 × 2 μm on the fourth side to retain a connection between the block and the grid bar. Then we polished the tip of the nanomanipulator needle and its attachment site on the block with 1.5 nA FIB current, and welded the block to the tip of the needle before releasing it completely from the grid. The welding conditions were as follows: 16 polishing patterns, 0.3 × 2 μm each, milling direction from the needle toward the copper block, FIB current 1.5 nA, calculated depth 1 μm. The patterns were positioned at the edge of the block-needle contact site, with 90% of the patterns’ length located over the copper block. After this, the sample holder was unloaded, and we prepared the microscope for the cryo-workflow.

Once a region of interest was located within the frozen organoids sample in the HPF carrier by means of cryo-FLM and following cryo-vEM, the area was targeted for the preparation of a sample block with dimensions of 45 × 25 μm. Large trenches were milled around the block at a stage tilt of 54°, FIB current of 7 nA, and a depth of 50 μm, to allow access for the nanomanipulator: 60 × 100 μm behind the block, and 10 × 40 μm and 10 × 20 μm on the sides to retain the connection to the bulk HPF specimen on the side of the shorter pattern. A 60 × 100 μm trench in front of the block was opened already during preparation for the cryo-vEM. Once the sample block dimensions were defined, the stage was tilted to −4° to perform an undercut (FIB milling current 7 nA) and release the block from the bottom side. The stage was tilted back to 54° and the nanomanipulator needle inserted. The nanomanipulator was pre-cooled to −185 °C (stabilization time after the external Dewar filling: 1 hour; recommended re-filling frequency is every hour to ensure optimal thermal and mechanical stability). The needle was brought into contact with the polished top of the target sample block (as viewed from the FIB perspective) and welded via the copper block using the following welding conditions: 18 polishing patterns, 0.3 × 4 μm each, milling direction from the sample block toward the copper block, FIB current 1.5 nA. Once the attachment was established, the remaining connection between the sample block and the HPF specimen was removed (3 nA, depth 50 μm), and the block was moved to the receiving lift-out grid by means of the nanomanipulator.

Before deposition onto the receiving grid, the stage was tilted to 14° to create a shallow angle of 10° between the FIB (positioned at 54° with respect to the SEM column) and the grid surface. Then, the sample block was lowered onto the grid such that the edges of the block rested on the parallel grid bars. The edges were welded to the grid bars via copper redeposition using the same parameters as for attaching the sample block to the copper block, with 4-5 patterns on each side. Finally, the bottom 5 μm of the block was milled with a rectangular pattern of 50 × 1 μm, current of 1.5 nA, depth of 30 μm, to generate a lamella. The width of the milling rectangle was set slightly broader than that of the block to mitigate re-attachment of the lamella to the block of the remaining lift-out volume due to material redeposition. This procedure was repeated to generate additional lamellae on the receiving grid. After the last lamella was detached from the nanomanipulator, the latter was retracted into the parking position, and the cooling Dewar refilling was stopped. We proceeded with step-wise thinning of the resulting lamellae to the final target thickness of ~200 nm. First, milling from 5 μm to 3 μm thickness was achieved with consecutive top and bottom polishing rectangles, with the milling direction towards the lamella, using a FIB current of 1.5 nA. Next, the lamellae were thinned to 1 μm using similar patterns with 700 pA, to 0.5 μm with 300 pA, and the final target thickness of 200 nm with 50 pA.

The sample holder was unloaded from the Zeiss Crossbeam 550 under cryo-conditions, the HPF sample and the lift-out grid were stored in sealed grid boxes under LN_2_ for future use.

### Cryo-ET data acquisition and processing

Lamellae were loaded into a Titan Krios G4 with 300 keV cold FEG, Selectris X energy filter, and Falcon 4i DED controlled by SerialEM Version 4.1.6 64-bit^60^. Intermediate-magnification (11.09 Å/pixel) montage overviews of the lamellae were acquired at a defocus of −70 μm. Areas for tilt-series data acquisition were selected to target prominent cellular features and cell-cell interfaces. Tilt-series acquisition was done at a pixel spacing of 3.03 Å/pixel, using a dose-symmetric scheme^61^ with 2° increments, a tilt range of 60° to −60°, a defocus range of 1.5 to 3.5, and an exposure dose of 2.7 e^−^/Å^2^ per tilt. Raw movies acquired in EER format^64^ were converted to TIFF format using RELION 3.1^65^. Movie-frame motion correction and contrast transfer function (CTF) estimation were performed with WarpTools^66^, after which the corrected tilt images were assembled into tilt-series stacks. Each tilt series was screened using an in-house developed script, and failed tilt images were excluded from further processing. Initial tilt-series alignment was performed with AreTomo^67^ through the WarpTools workflow. The resulting alignments were subsequently refined using MissAlignment^68^. Following model training,the optimized alignment parameters were applied to the corresponding tilt series. Tomograms were then reconstructed using WarpTools. For denoising, even- and odd-frame half-datasets were generated with WarpTools and used to reconstruct paired half-tomograms. Masks restricting the analysis to the lamella volume were generated using the Etomo interface of IMOD 4.11.12^63^ and Slabify^69^. The masked half-tomograms were subsequently processed with IsoNet2^70,71^ for denoising.

All schematics were created with BioRender (BioRender.com) and Adobe Illustrator.

## Acknowledgments

We are grateful for the support from the EMBL electron microscopy core facility, cryo-EM platform and IT Services. In particular, we thank Dr. Anna Steyer from the EMBL Imaging Center for expert help with cryo-vEM data acquisition, and Dr. Martin Fritsch from Leica Microsystems for invaluable support with LASX software usage. We acknowledge the in-kind support of Oxford Instruments through the provision of an OmniProbe 350 cryo system on loan and the associated technical expertise, collaboration, and support provided by Kim Larsen, John Lindsay, and Louise Hughes. E.D. was supported by a fellowship from the EMBL Interdisciplinary postdoctoral program (EI4POD) under Marie Skłodowska-Curie Actions COFUND (847543). J.M. acknowledges support from the EMBL and a grant within the iNext Discovery framework (871037).

